# Root Colonization and Growth Promotion of Soybean, Wheat and Chinese Cabbage by *Bacillus cereus* YL6

**DOI:** 10.1101/353946

**Authors:** Yongli Ku, Guoyi Xu, Fawei Wang, Haijin Liu, Xiangna Yang, Xiaohong Tian, Cuiling Cao

## Abstract

Phosphate-solubilizing bacteria (PSB) have been isolated and used in agricultural production. However, comprehensive research on PSB colonizing the rhizosphere of different plants and promoting plant growth is lacking. This study was conducted to study the growth-promoting effects and colonizing capacity of the PSB strain YL6. The YL6 strain not only increased the biomass of pot-planted soybean and wheat but also increased the yield and growth of Chinese cabbage under field conditions. The promotion of growth in these crops by strain YL6 was related to its capacities to dissolve inorganic and organic phosphorus and to produce a certain amount of indole-3-acetic (IAA) and gibberellin (GA). After YL6 was applied to soybean, wheat and Chinese cabbage, the rhizosphere soil available phosphorus (available P) content increased by 120.16%, 62.47% and 7.21%, respectively, and the plant total phosphorus increased by 198.60%, 6.20% and 78.89%, respectively, compared with those of plants without the addition of YL6. To determine whether the phosphate solubilizing bacteria colonized these plants, YL6 labeled with green fluorescent protein (YL6-GFP) was inoculated into plant rhizospheres. YL6-GFP first colonized the root surface and hairs and then penetrated into intercellular spaces and vessels. Collectively, these results demonstrate that YL6 promoted the growth of three different crops and colonized them in a similar way and therefore provide a solid foundation for probing into mechanisms by which phosphate-solubilizing bacteria affect plant growth.

## Introduction

Phosphorus as one essential element and the primary growth-limiting factor for plant growth plays significant roles in some major metabolic processes, such as photosynthesis, respiration, and molecular synthesis [1]. Phosphorus as a nonrenewable resource has attracted much attention [2]. In soil, the primary phosphorus forms are apatite, calcium phosphate and organic phosphorus, which have relatively low phosphorus availabilities for plants [3]. Readily available P deficiencies restrict crop yields. To address this problem, large amounts of phosphate fertilizers must be applied. Of phosphorous fertilizer, 80-90% is fixed as insoluble P and only a very small part is available for plants [4]. The soil accumulation of insoluble P leads to environmental pollution [5]. Thus, approaches to increase efficiencies of phosphate fertilizers and make soil insoluble P environmentally friendly are urgently required.

Phosphate-solubilizing bacteria (PSB) are required for a series of biochemical reactions to convert insoluble P to absorbable and available forms [6,7]. The primary mechanisms behind these reactions are the following: (a) PSB secret organic acids, such as gluconic, acetic, and citric acids, to dissolve mineral complexes; and (b) PSB produce phosphatase enzymes to degrade insoluble organic P [8,9]. Similar to other plant growth-promoting rhizobacteria (PGPR), some PSB strains can promote plant growth by producing indole-3-acetic (IAA) and gibberellin (GA) [10,11]. Therefore, application of PSB in agricultural production is an important way to achieve phosphorus cycling and sustainable development in farmlands. Many studies demonstrate that application of PSB into the plant rhizosphere can improve plant growth [12]. For example, Walpola and Yoon [13] verified promoting effects of PSB on tomato growth.

Studying the colonization and distribution of endophytic bacteria in plants enriches our knowledge about how bacteria affect plant growth. The green fluorescent protein (GFP), first isolated from *Aequorea victoria*, is widely used as a marker in studying gene expression and bacteria localization [14,15]. Furthermore, GFP-labeled bacteria are easily detected without requiring isolation, culturing and identification; thus, GFP is an ideal tool for studying the colonization of endophytic bacteria in plants. Actually, many PGPR tagged with GFP are employed to study microbial colonization patterns, leading to certain advances [16–18].

However, because of a lack of systematic studies, whether PSB strains can promote the growth of and colonize in different crops in a similar way remains unclear. In this study, the YL6 strain, a bacterial strain that dissolves inorganic and organic phosphorus, was applied to soybean (*Glycine max* L. Merr.), wheat (*Triticum aestivum* L.) and Chinese cabbage (*Brassica rapa* L., *Chinensis Group*) under different conditions. Additionally, YL6-GFP, a GFP-marked YL6 strain, was employed to study the PSB colonization of the three different crops. The YL6 strain similarly promoted the growth of these plants by dissolving inorganic and organic phosphorus and producing certain amounts of IAA and GA in soil. Moreover, YL6-GFP also colonized the different plants in a similar approach.

## Materials and methods

### Ethics statement

This work was conducted in our scientific research field for PSB *Bacillus cereus* studies, which is owned by our institution. Therefore, no specific permissions were required for these using these locations or performing the study. And this field did not involve endangered or protected species.

### Test strain

The PSB YL6 maintained by our laboratory was isolated from the rhizosphere of Chinese cabbage at a soil depth of 10 cm at the Yangling Experiment Farm of Northwest A&F University (34.30° N, 108.08° E), Shaanxi, China. Based on 16s rRNA testing, YL6 was identified as *Bacillus cereus* of which the GenBank accession number was KX580383 [9].

### Phosphate-solubilizing capacities and growth-promoting substances of YL6

Purified YL6 was inoculated into 100 ml of inorganic phosphorus liquid medium (glucose 10 g/l, (NH_4_)_2_ SO_4_ 0.5 g/l, NaCl 0.3 g/l, KCl 0.3 g/l, MgSO_4_•7H_2_O 0.3 g/l, FeSO_4_•7H_2_O 0.03 g/l, MnSO_4_•4H_2_O 0.03 g/l, CaCO_3_ 10 g/l, pH 7) [19] and incubated for 24, 48 and 72 h at 30°C on a shaker running at 180 rpm in triplicate to measure soluble P and organic acids in the culture medium. For each repetition, 20 ml of the medium was taken and centrifuged at 13,000 g for 10 min to obtain cell-free supernatants. The soluble P contents of the supernatants were determined by the molybdate blue method [20]. To determine organic acids by High-Performance Liquid Chromatography (HPLC), the supernatants were filtered with 0.45-μm nylon filters (Millipore Corp, Billerica, MA, USA). Then, 20 μL of this leachate was injected into the HPLC instrument (Essentia LC-15C, Japan), which was equipped with a C-18 column and run at the flow rate of 1 ml/min using 90:10 (v/v) methanol-phosphate buffer (10 mmol/l) of pH 2.7 as the mobile phase; this process was monitored at 210 nm [21–23]. Additionally, purified YL6 were inoculated into 100 ml of organic phosphorus liquid medium (Extractum carnis 5 g/l, peptone 10 g/L, NaCl 5 g/l, pH 7.0; after 20 min, the medium was sterilized at 121°C, and 40 ml of a mixture of yolk and physiological saline at the ratio of 1:1 was added) [24] in triplicate and incubated at 30°C for 24 h on a shaker running at 180 rpm. The production of soluble P and acid, alkaline and neutral phosphatase activity of YL6 were measured by the p-nitrophenyl phosphate method [25].

Purified YL6 were inoculated into 100 ml of LB liquid medium in triplicate and incubated at 30°C for 24 h on a shaker running at 180 rpm. For each repeat, 20 ml was centrifuged at 13,000 g for 10 min to obtain cell-free supernatants, which were used to measure the YL6 capacity to produce IAA and GA by the Salkowski colorimetric assay [26] and fluorimetry assay [27].

### Influences of YL6 on rhizosphere soil and biomass of different crops Soybean pot experiment

Our laboratory purchased soybean seeds (Zhonghuang 13). Calcareous soil that was not planted and fertilized over many years was collected from the Medicinal Garden of Northwest A&F University (34.25° N, 108.06° E) and then air-dried and sifted through a 2 mm sieve. The bottom diameters and heights of the test pots were 28 cm. Each pot was filled with 3 kg of soil, and 6 soybean seeds were sown in each pot. The YL6 strain was incubated at 30°C×180 rpm for 48 h in LB liquid medium to produce YL6 agent. The three treatments were the following: (1) YL6 (180 ml of 2.8×10^8^ cfu/ml YL6 agent added at soybean seedling emergence, (2) M (Pure medium with no strains added at the same time as the YL6 treatment), and (3) CK (the control without any treatments). All the treatments were repeated five times. After seedling germination, two seedlings in good growth condition were kept in each pot. The soil absolute water content was maintained at 20-22%. When the soybean seedlings had three compound leaves, 8 plants were randomly collected in each of the treatments.

Root and shoot fresh and dry weights and total fresh and dry biomass of soybeans were measured directly. Dry weights were determined after drying for 48 h at 80°C in an air oven, and the R/S ratio was the ratio of root dry weight and shoot dry weight [28]. The soybean vegetative growth and bean pod indexes (including stem diameter, primary branch number, and pod length, width and number) were directly measured.

Each fresh soil sample (2 g) collected from the soybean rhizosphere was placed in a triangular bottle filled with 18 ml of sterile water and shaken at 30°C on a shaker running at 180 rpm for 30 min. Then, this mixture was left to settle for a time. Suspensions (1 ml) taken from mixture were diluted to 10^−5^ and uniformly smeared on inorganic phosphorus agar medium by the plate-smearing method [29]. The smeared agar medium was incubated (in darkness at 30°C) in a constant temperature incubator for 3 days. At the end of the incubation, the colony numbers of PSB in the soil samples were tallied (cfu/g). The soil available P was extracted by the bicarbonate method and measured by the molybdate blue method [30]. Dried individual plants were ground and sifted through a 1 mm sieve, treated with H_2_SO_4_-HClO_4_ and the phosphorus contents were measured by the vanadium molybdenum yellow method [31].

### Wheat pot experiment

The seeds of wheat (Xiaoyan 22) were from storage in our laboratory in this experiment. The soil and pots were the same as those in the soybean experiment. Each pot was filled with 3 kg of soil, and 20 wheat seeds were sown in each pot. The 3 treatments in the experiment were as follow: (1) YL6 (180 ml of 2.8×10^8^ cfu/ml YL6 agent added at wheat seedling emergence), (2) M (Pure medium with no strains added at the same time as the YL6 treatment), and (3) CK (the control without any treatments). After wheat seedlings had 3 leaves, 15 plants were kept in each pot in good growth condition. Each treatment was repeated 3 times. The soil absolute water contents were maintained at 20-22%. At the tillering stage, 36 wheat plants growing identically were randomly selected from 3 pots of each treatment.

The height, root length, root fresh and dry weights, shoot fresh and dry weights and total fresh and dry plant biomass of different treatments were measured directly. The same methods as in the previous experiment were used to determine soil available P, plant phosphorus and the number of phosphate-solubilizing rhizosphere bacteria.

### Field experiment for Chinese cabbage

Seeds of Chinese cabbage (Shanghaiqing) were from storage in our laboratory. From August 29 to October 10, 2016, the experiment was conducted at the Yangling Experiment Farm of Northwest A&F University (34.30° N, 108.08° E). The soil was calcareous soil that was kept idle over many years. The three treatments were the following: (1) YL6 (1500 ml of 2.8×10^8^ cfu/ml YL6 agent added at cabbage seedling emergence), (2) M (Pure medium with no strains added at the same time as the YL6 treatment), and (3) CK (the control without any treatments). The experimental fields were divided into 15 identical plots (2 m×2 m). The design was a randomized block with three replicates. In the growing season of Chinese cabbage, the experimental farmland was managed scientifically. Then, we sampled ripe vegetables randomly with the same growth condition for further analysis. Root fresh and dry weights, shoot fresh and dry weights and total fresh and dry plant biomass of different treatments were measured directly. To measure the quality of Chinese cabbage, vitamin C was determined by the molybdenum blue colorimetric method, cellulose and soluble sugar were determined by the anthrone colorimetric method, soluble protein was determined by the Coomassie blue staining method, and nitrate nitrogen content was measured by nitration of salicylic acid colorimetry [32].

### YL6 colonization on root surfaces of soybean, wheat and Chinese cabbage seedlings

GFP-labeled YL6 was constructed in our laboratory, and the stability was tested [9]. Soybean, wheat and Chinese cabbage were cultured by the sandy culture method. The sand (passed through a 24 mesh sieve) was washed clean with tap water and sterilized at 120 °C in an oven for 48 h. Then, 1 kg of sterilized sand was put into individual boxes (195 mm×146 mm×65 mm). Tap water was added to the pot to maintain absolutely the water content range within 8-12%. The seeds of soybean, wheat and Chinese cabbage were sterilized with 10 mL of 2% NaClO (13% active Cl^−^ content) for 10 min. Then, the seeds were rinsed 5 times with sterile water. The aseptic seeds were soaked in warm water for 30 min, and those that floated up on the surface of the water were removed. The remaining seeds were evenly sown on three stacked wet filter papers in culture dishes. The culture dishes were incubated at 28 °C in the darkness in an incubator for 2 days. Water was added every 8 h to keep filter papers wet. After germination, seeds were incubated at 28 °C under the day/dark pattern of 16 h/8 h. Subsequently, the different seeds were transplanted into different culture boxes with sand. When the seedlings grew 3 primary leaves and developed a root system, the YL6-GFP bacterial suspension (2.8×10^8^ cfu/ml) diluted twice (1.35×10^8^ cfu/ml) was added to the culture boxes. At 3, 6 and 9 days after the addition, the seedlings were collected and rinsed with sterile water. Then, some root tissues were cut into tiny pieces in a crisscrossed pattern with a sterilized razor blade for bacterial colony observations, with the pieces then placed in glass dishes by the hydrostatic tablet compressing method. The YL6-GFP endogeny colonization on fresh roots was visualized by fluorescence microscopy (Olympus CCD-DP26).

### Statistical analyses

Microsoft Excel 2007 was employed to process the data. The comparisons among the treatments were performed using the least significant difference (LSD) at *P*<0.05. Adobe Photoshop CS6.0 was used for photo combinations.

## Results

### Phosphate-solubilizing capacities and production of growth-promoting substances of YL6

Because of the difference between the mechanisms by which YL6 dissolved inorganic and organic phosphorus, the solubilizing phosphorus capacities of the YL6 strain in different liquid culture media were further investigated (Table 1). First, YL6 was inoculated into organic phosphorus liquid medium to determine the activity of acid phosphatase, alkaline phosphatase and neutral phosphatase and the accumulation of available P at the different culture times. When the activities of these phosphatases were the highest, the content of available P in the medium was also the highest. Thus, the concentration of available P increased to 152.253 μg/ml at 48 h of culture. Additionally, YL6 secreted oxalic, malonic and succinic acids to increase available P in the inorganic phosphorus liquid medium (Table 1). Furthermore, other growth-promoting substances were also detected. For example, the concentrations of IAA and GA increased to 29.503 and 30.615 mg/L, respectively, at 24 h of incubation (Table 1). These results indicated that the YL6 strain had strong capacities to dissolve different forms of phosphorus into available P for plants.

**Table 1.**
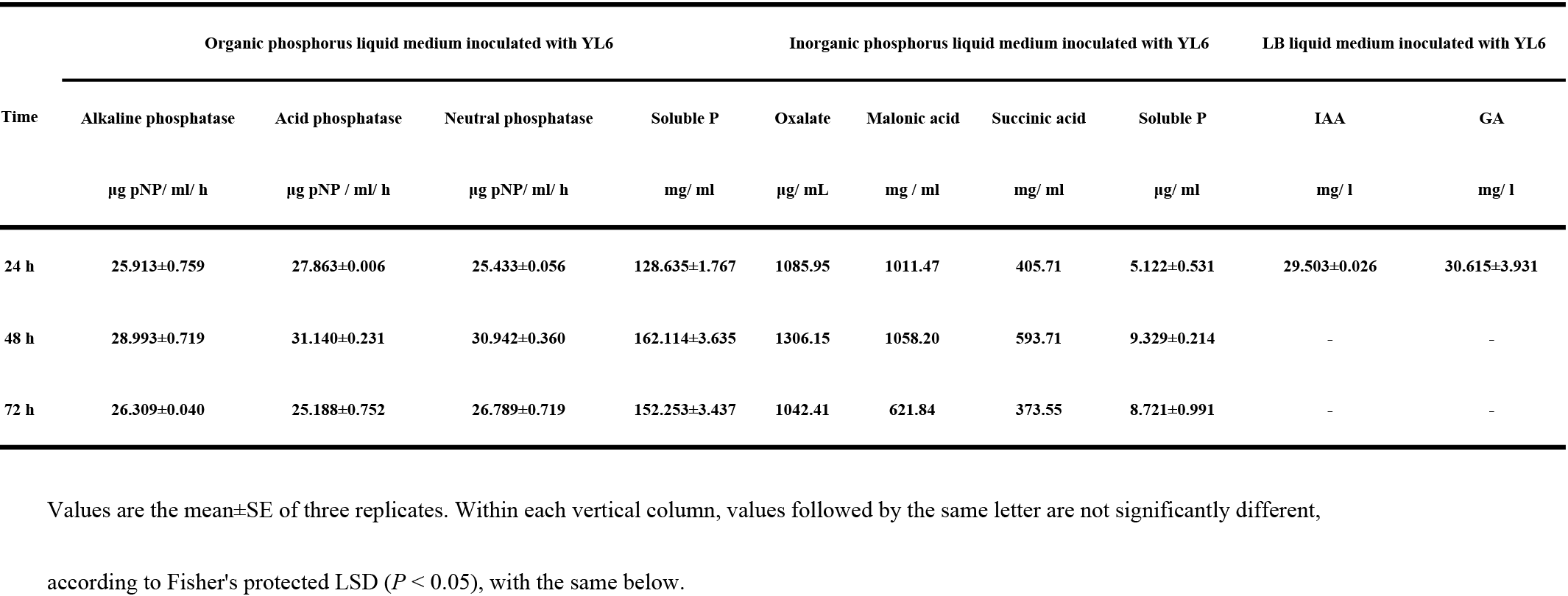
The ability of YL6 to dissolve phosphorus and produce auxins.

### Influence of YL6 on biomass and quality of the different crops

To test whether this PSB could promote plant growth, soybean, wheat and Chinese cabbage carrying the YL6 strain were planted. Based on the phenotypes of these crops, YL6 treatment obviously improved growth (Figs 1-3). Furthermore, the fresh and dry weights of roots, shoots and plants of the crops treated with the YL6 strain were much higher than those of the M and CK groups (Table 2). For soybean, the R/S ratio of the YL6 group increased by 28.8% (*P* < 0.05) compared with that of the CK group. Different indexes of soybean growth and the bean pod (stem diameter, primary branch number, and pod length, width and number) also increased significantly with YL6 strain inoculation compared with those of the control (Table 3), and the number, length and width of soybean pods increased significantly by 16.81%, 94.34% and 251.13%, respectively (*P* < 0.05). YL6 increased wheat and Chinese cabbage biomass compared with that of CK and M groups. In addition to the biomass indexes, the nutritional values of Chinese cabbage were also determined. We found that leaf vitamin C, soluble sugar, soluble protein and cellulose were the highest in the YL6 strain treatment, reaching 2.105, 15.610, 15.695 and 42.539 mg/g (*P*<0.05), respectively (Table 4). Another important index of Chinese cabbage is the content of nitrate nitrogen, which can be transformed into nitrates harmful to humans. Although the nitrate nitrogen contents in these groups were not different from one another (*P*<0.05), the nitrate nitrogen content in the YL6 group was lower (Table 4). Based on these results, the PSB YL6 strain significantly improved the growth of the different crops.

**Fig 1.**
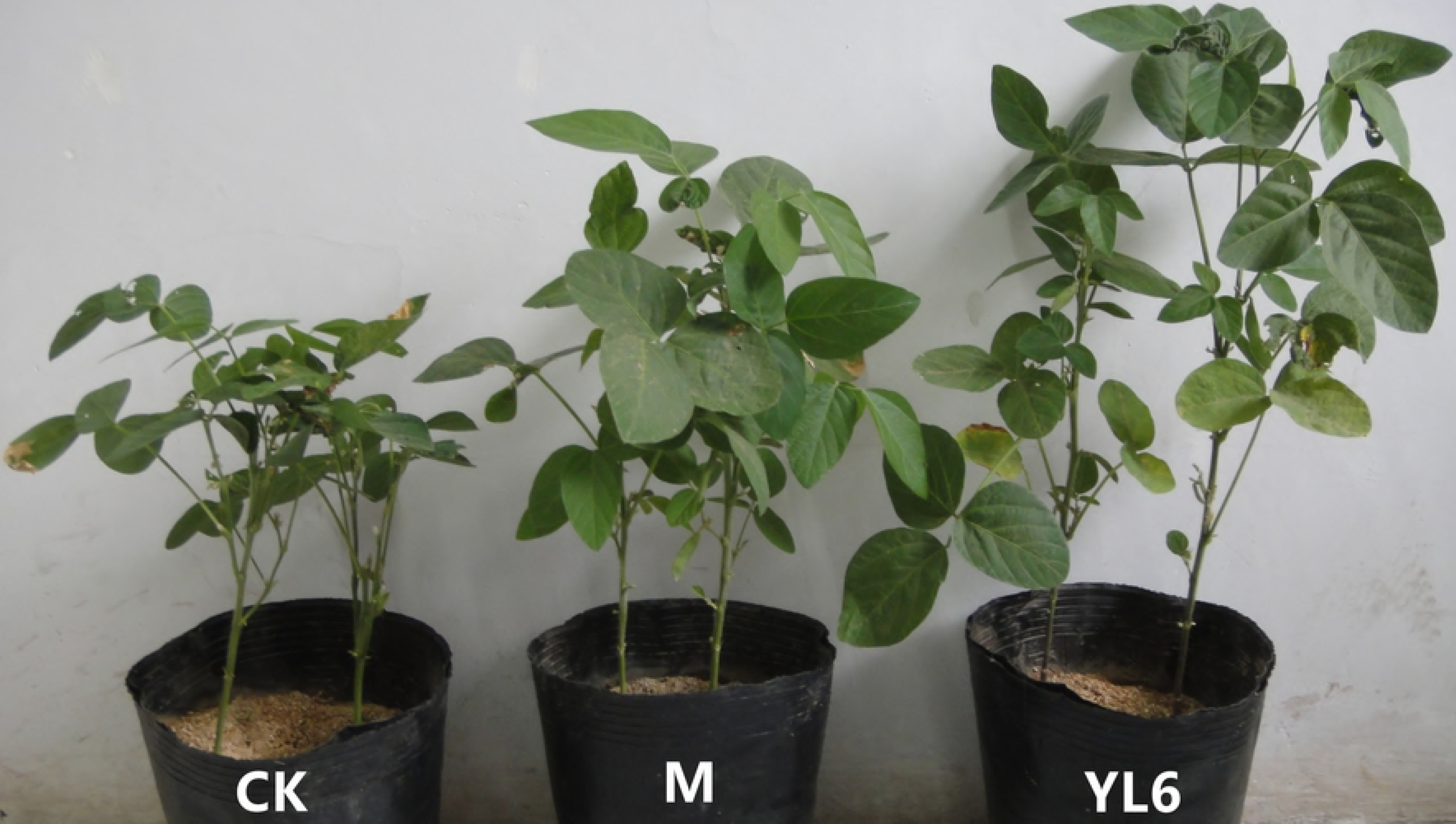
Effect of YL6 on promotion of growth in soybean. **CK**, control group; **M**, treatment with liquid medium without the strain; **YL6**, treatment with the YL6 agent.

**Fig 2.**
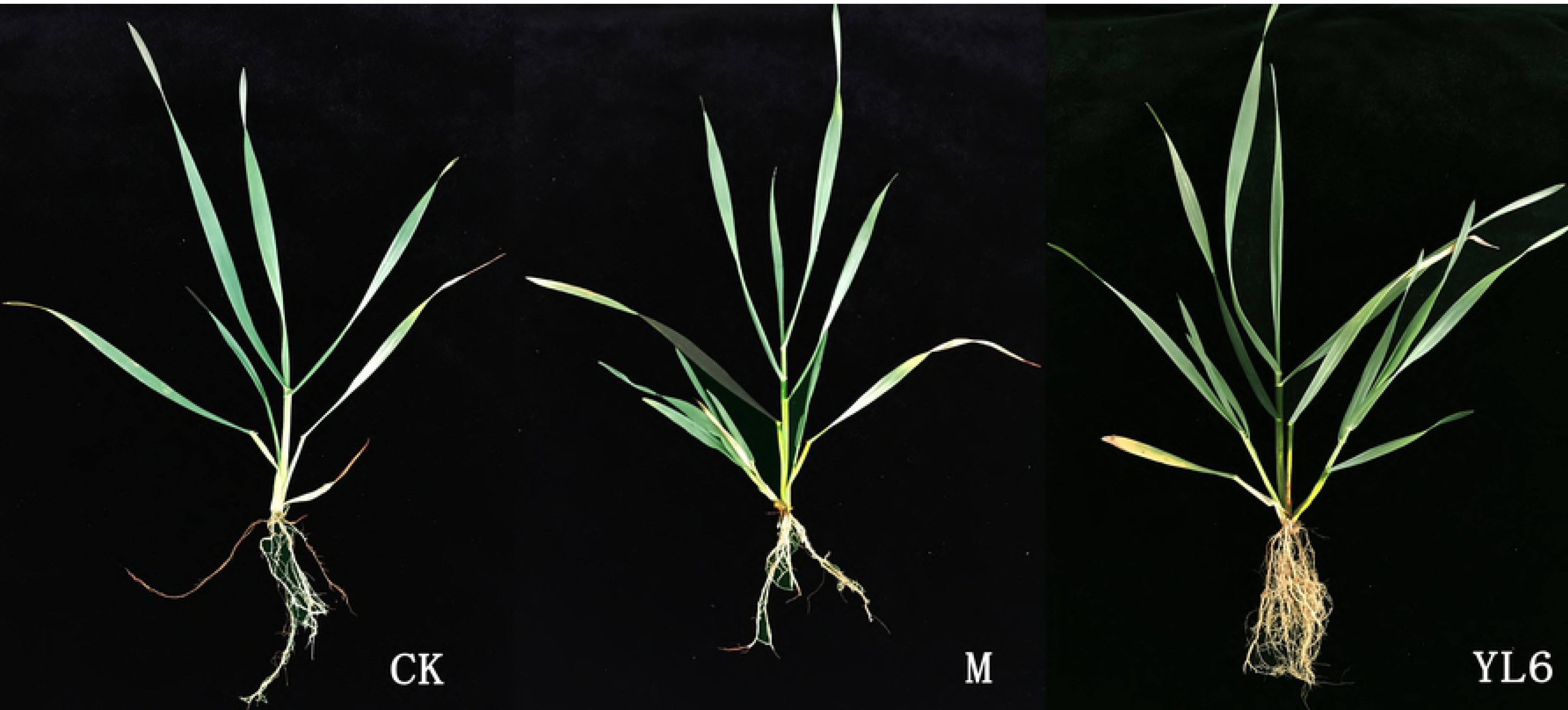
Effect of YL6 on promotion of growth in wheat. **CK**, control group; **M**, treatment with liquid medium without the strain; **YL6,** treatment with the YL6 agent.

**Fig 3.**
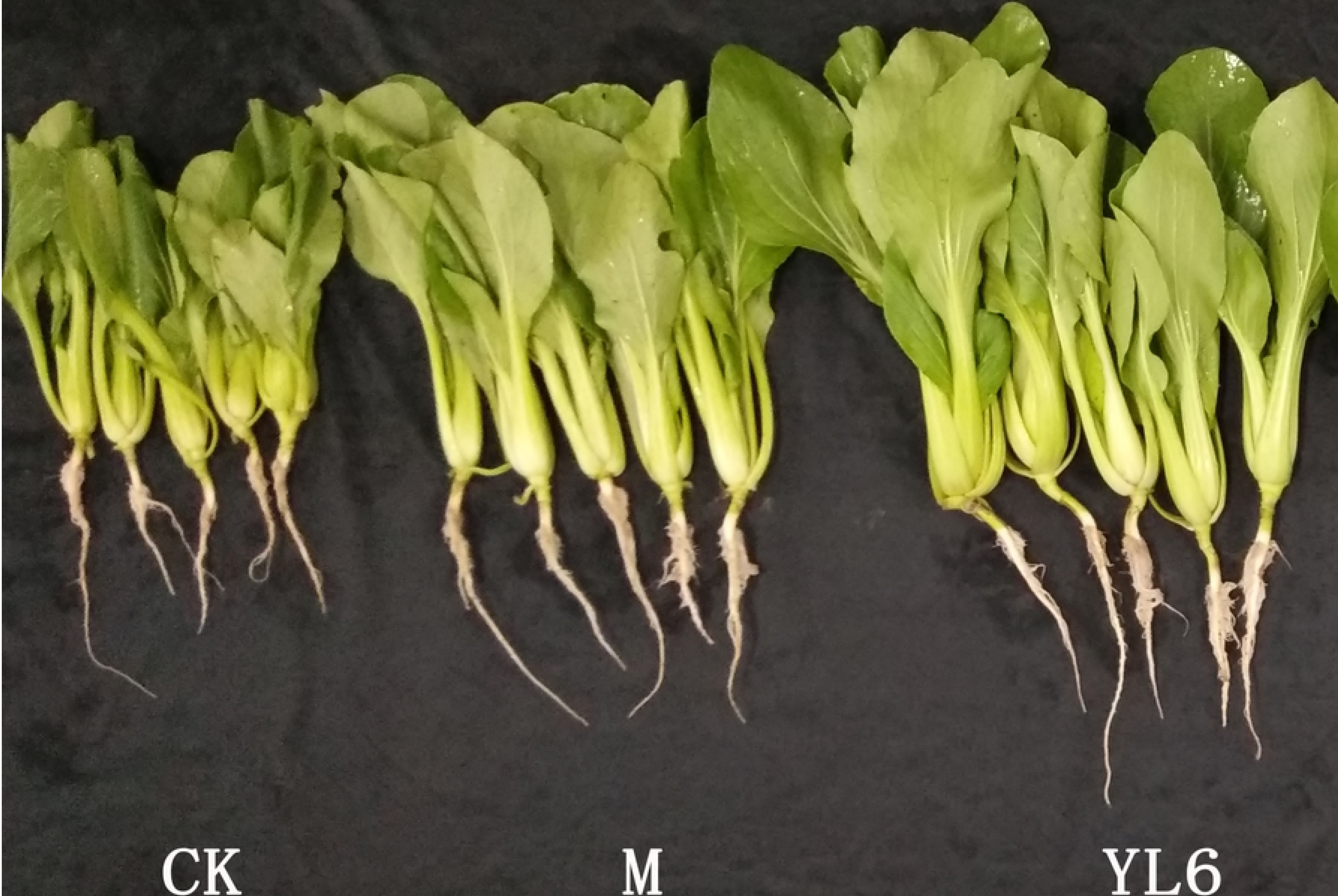
Effect of YL6 on promotion of growth in Chinese cabbage. **CK**, control group; **M**, treatment with liquid medium without the strain; **YL6,** treatment with the YL6 agent.

**Table 2.** Effect of YL6 on soybean, wheat and Chinese cabbage biomass.

| Crop type | Treatment | Shoot fresh | Root fresh | Shoot dry | Root dry | Fresh | Dry | R/S<br>(DW/DW) | Yield/plot<br>kg |
| --- | --- | --- | --- | --- | --- | --- | --- | --- | --- |
|  |  | weight | weight | weight | weight | biomass | biomass |  |  |
|  |  | g/plant<sup>-1</sup> | g/plant<sup>-1</sup> | g/plant<sup>-1</sup> | g/plant<sup>-1</sup> | g/plant<sup>-1</sup> | g/plant |  |  |
| Soybean | CK | 2.72±0.06c | 1.36±0.06c | 0.42±0.02c | 0.10±0.04c | 4.08±0.15c | 0.52±0.01c | 0.25±0.02b | - |
|  | M | 3.47±0.19b | 1.52±0.05b | 0.61±0.00b | 0.20±0.01b | 4.99±0.057b | 0.81±0.01b | 0.32±0.01a | - |
|  | YL6 | 4.65±0.10a | 2.75±0.07a | 0.74±0.02a | 0.24±0.01a | 7.40±0.14a | 0.97±0.02a | 0.32±0.01a | - |
| Wheat | CK | 0.55±0.18c | 0.04±0.00b | 0.14±0.02c | 0.02±0.01c | 0.65±0.26b | 0.16±0.03c | 0.16±0.01a | - |
|  | M | 1.13±0.16b | 0.09±0.01b | 0.27±0.01b | 0.04±0.01b | 1.15±0.04a | 0.30±0.02b | 0.13±0.01b | - |
|  | YL6 | 2.32±0.20a | 0.25±0.07a | 0.57±0.00a | 0.09±0.00a | 2.66±0.15a | 0.67±0.00a | 0.16±0.00a | - |
| Chinese cabbage | CK | 12.68±0.63d | 0.47±0.06d | 0.99±0.06d | 0.08±0.02d | 13.15±0.64b | 1.07±0.07c | 0.08±0.00b | 2.21±0.09c |
|  | M | 22.93±1.01c | 1.34±0.03c | 1.66±0.18c | 0.19±0.02c | 24.07±0.96b | 1.11±1.03c | 0.11±0.02a | 2.57±0.03c |
|  | YL6 | 43.61±5.62b | 1.57±0.23b | 2.74±0.29b | 0.25±0.05b | 44.76±5.33b | 3.04±0.24b | 0.11±0.02a | 3.87±0.39b |

**Table 3.**
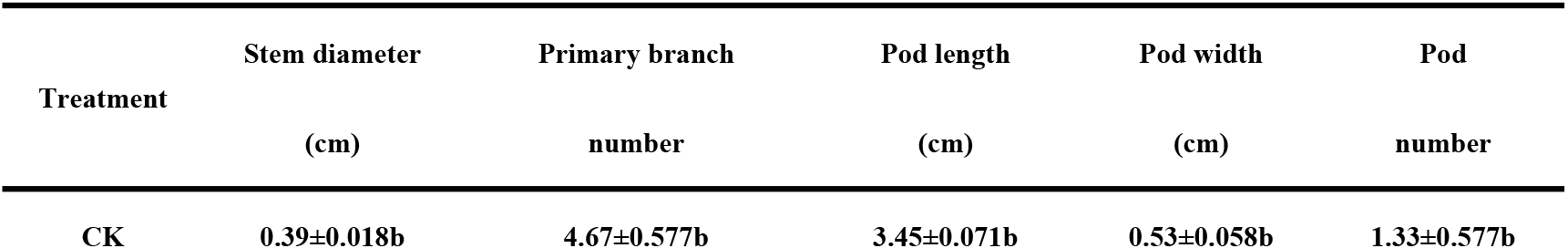
Effect of phosphate-solubilizing bacteria on soybean vegetative growth and bean pods.

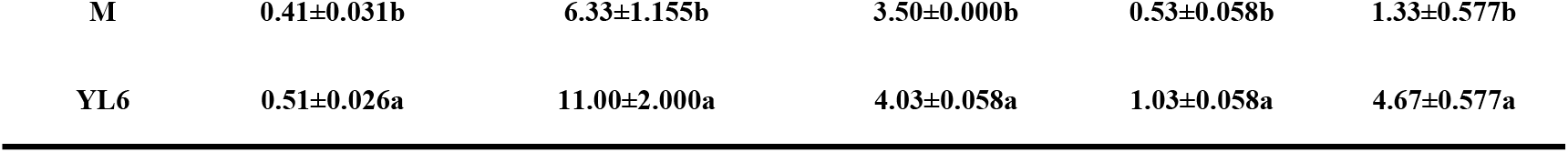

**Table 4.** Effect of different treatments on the quality of Chinese cabbage.

| Treatment | Cellulose<br>mg/g | Vitamin C<br>mg/g | Soluble sugar of<br>leaves<br>mg/g | Soluble protein of leaves<br>mg/g | Nitrate nitrogen of<br>leaves<br>mg/g |
| --- | --- | --- | --- | --- | --- |
| CK | 6.445±0.337c | 1.986±0.079b | 12.004±1.246c | 19.817±0.469c | 1.140±0.116a |
| M | 7.575±0.569c | 2.059±0.140b | 13.305±0.831c | 31.108±1.713b | 1.064±0.069a |
| YL6 | 15.610±2.613b | 2.105±0.127b | 15.695±0.672b | 42.539±4.491a | 1.062±0.052a |

### Plant phosphorus, number of soil phosphate-solubilizing bacteria and soil available P

The amount of plant phosphorus was determined to check whether the YL6 strain promoted soybean, wheat and Chinese cabbage growth by helping plants absorb soil phosphorus. First, PSB strains in rhizosphere soils of these crops were counted. Numbers of PSB strains increased with the application of YL6 and were 13, 35 and 10-fold those of the CK group in soybean, wheat and Chinese cabbage, respectively (Table 5). The increase in PSB strains generated more available P in soil (Table 6). For example, the soil available P content in the YL6 group was 120.16% higher than that in the CK group in the soybean pot experiment. The increase in available soil P was easily utilized by the plants, which led to an increase in the total P contents of soybean, wheat and Chinese cabbage (Table 7). The total P contents of soybean, wheat and Chinese cabbage in the YL6 group were 198.60, 6.20 and 78.89% higher than those in the CK groups, respectively. These results indicated that the YL6 strain similarly promoted the growth of different crops by increasing available soil P for plants.

**Table 5.** Effect of YL6 on the number of phosphate-solubilizing bacteria in rhizosphere soil (1×10^5^ cfu)

| Crop | CK | M | YL6 |
| --- | --- | --- | --- |
| Soybean | 0.009c | 0.08b | 0.12a |
| Wheat | 0.01c | 0.16b | 0.35a |
| Chinese cabbage | 0.35c | 1.80b | 3.50a |

**Table 6.** Effect of YL6 on soil available phosphorus (mg/kg)

| Crop | CK | M | YL6 |
| --- | --- | --- | --- |
| Soybean | 2.530±0.050b | 2.720±0.030b | 5.570±0.230a |
| Wheat | 13.788±0.654c | 18.942±1.332b | 22.401±1.378a |
| Chinese cabbage | 18.898±1.976b | 19.254±1.793b | 20.261±1.744a |

**Table 7.** Effect of YL6 on plant phosphorus (mg/g)

| Crop | CK | M | YL6 |
| --- | --- | --- | --- |
| Soybean | 1.220±0.050b | 2.081±0.030b | 3.643±0.230a |
| Wheat | 1.579±0.016b | 1.671±0.030a | 1.677±0.032a |
| Chinese cabbage | 1.179±0.212b | 1.395±0.000b | 2.109±0.130a |

### Root colonization of the different crops by YL6

In addition to testing the capacities of the PSB to promote the growth of crops, the colonization by a GFP-labeled strain of PSB, YL6-GFP, was examined. Under fluorescence microscopy, the YL6-GFP strain was detected only at root hairs of the three crops three days after inoculation, suggesting that YL6-GFP first attached to root hair surfaces of the crops and then penetrated into the root hairs (Fig 4A and 4C; Fig 5A and 5C; Fig 6A and 6C). Many fluorescent points were observed in the intercellular spaces of the root cortex of soybean on the sixth day after inoculation (Fig 4E and 4G). YL6-GFP was also distributed in the intercellular spaces of and even inside epidermal cells (Fig 5E, 5G, and 5I). Additionally, this strain colonized on the surfaces of epidermal cells of Chinese cabbage (Fig 6G). Finally, the YL6-GFP strain appeared in the vessels of these plants, because the green fluorescent strain was detected in the vessels of branch roots, roots and different samples of soybean, wheat and Chinese cabbage on the ninth day after inoculation (Fig 4I and K; Fig 5K; Fig 6I and K). These results suggested that the YL6-GFP colonized the different plants through a similar process: attaching first to the root hair, then penetrating into the vessels and finally expanding into the other organs.

**Fig 4.**
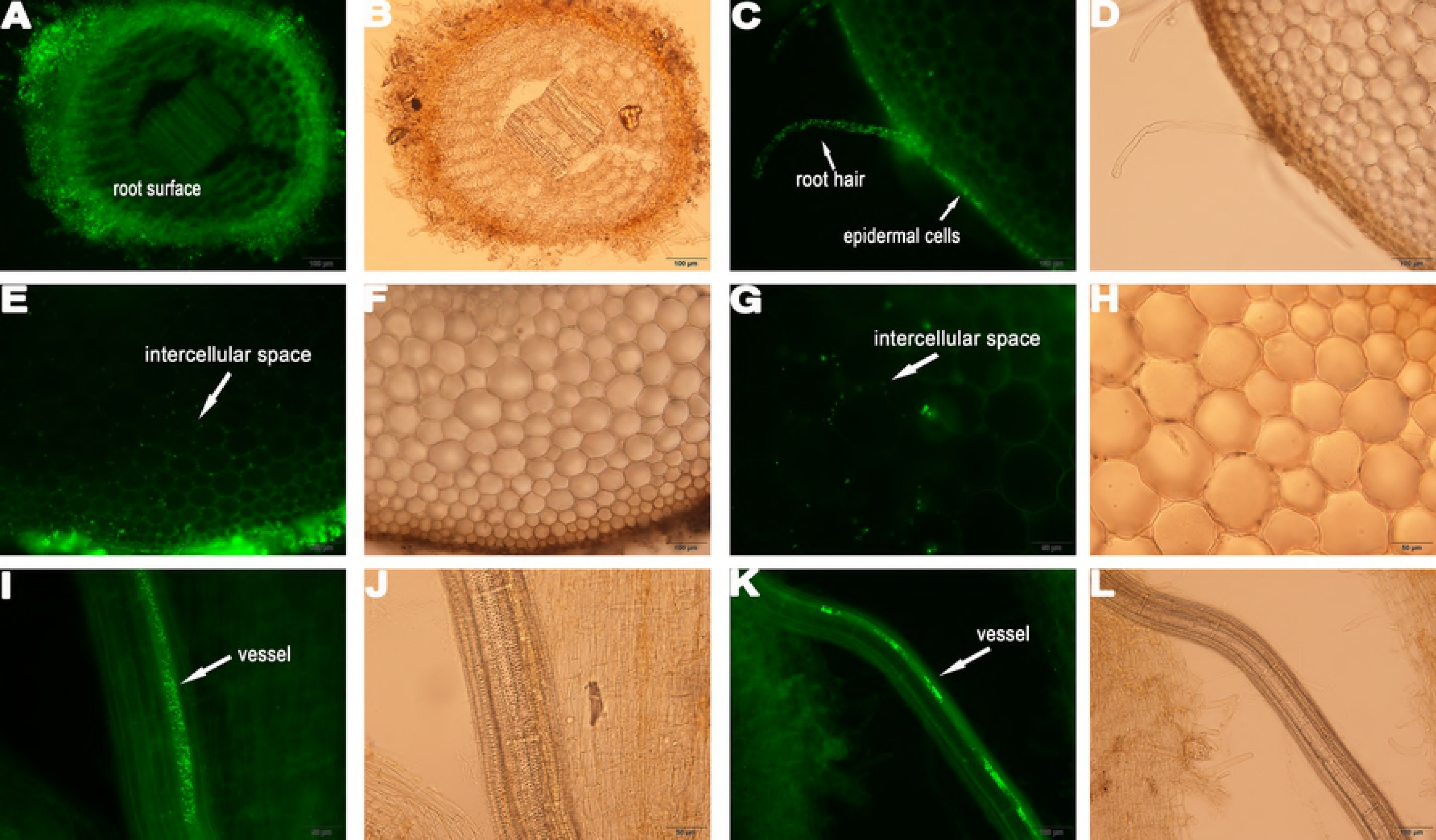
Colonization process of YL6-GFP in soybean root tissue. **A** longitudinal picture of root; **C** picture of root hairs and surface of primary root; **E**, **G** intercellular space of cortex; **I**, **L** YL6-GFP in the vessels of branch roots. **B**, **D**, **F**, **H**, **J**, and **L** were taken under bright field.

**Fig 5.**
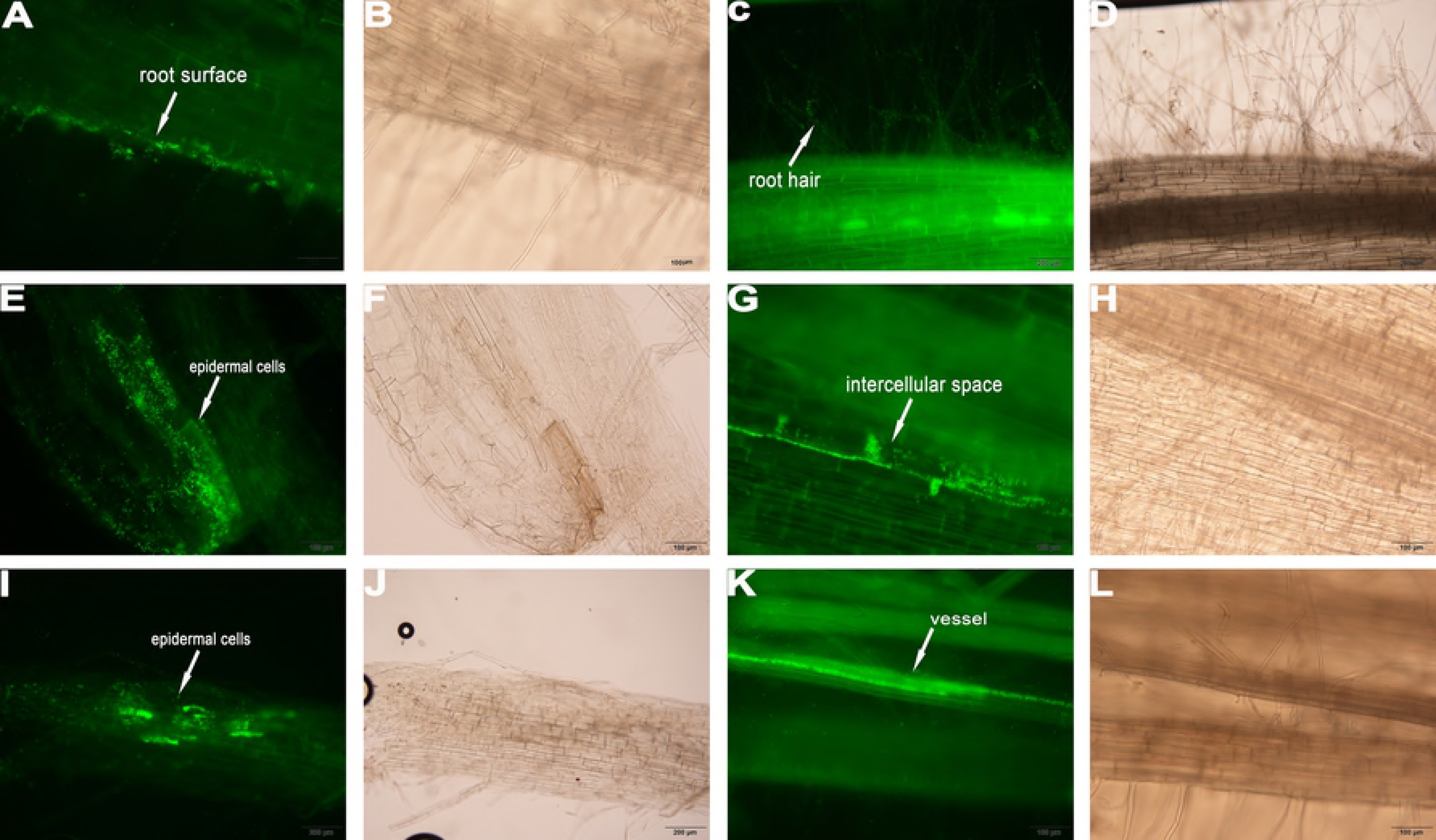
Colonization process of YL6-GFP in wheat root tissue. **A**, **C** surface of primary root and root hairs; **E**, **I** some epidermal cells colonized by YL6-GFP; **G** some bacteria distributed in intercellular spaces between cells; **K** YL6-GFP in the vessels of branch roots. **B**, **D**, **F**, **H**, **J**, and **L** were taken under bright field.

**Fig 6.**
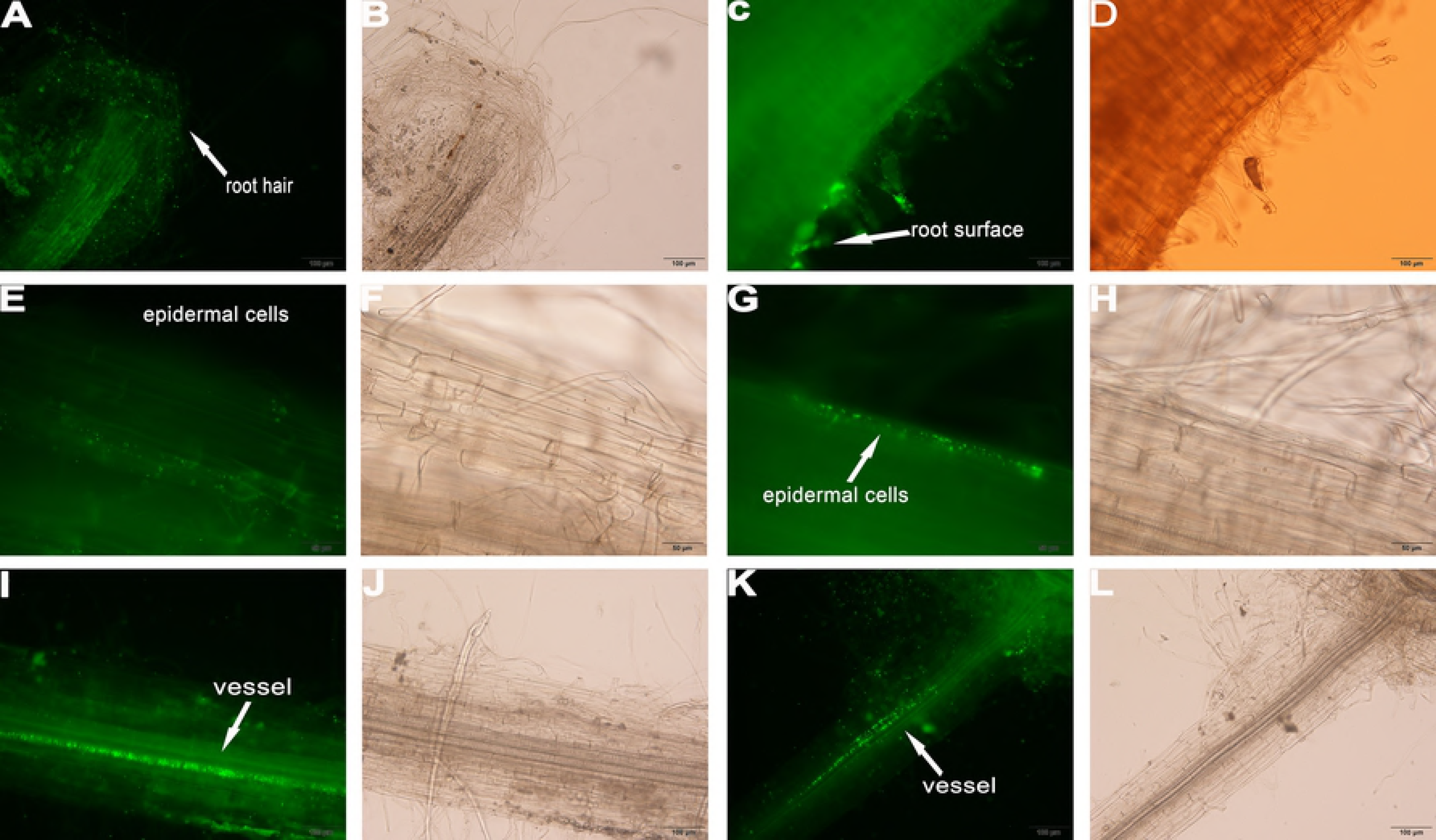
Colonization process of YL6-GFP in Chinese cabbage root tissue. **A**, **C** YL6-GFP on the root hairs and surface of primary root; **E**, **G** some epidermal cells colonized by YL6-GFP; **I**, **K** YL6-GFP in the vessels. **B**, **D**, **F**, **H**, **J**, and **L** were taken under bright field.

## Discussion

Phosphorus is a limiting factor in agricultural production [33]. Calcareous soil is widely distributed in northwest China, and lack of available P in the region restricts agricultural activities. Research on PSB provides new ideas to solve the problem of low phosphorus availability. According to many scholars, *Bacillus subtilis* KPS-1, *Pseudomonas dimnuta* RS-1, *Xanthomanas* sp. RS-3, *Exiguobacterium* sp. RS-4, and *Alcaligenes faecalis* Ss-2 have remarkable phosphorus-dissolving capacities [34–36]. Biological methods that use PSB to solve agricultural problems have huge potential in the future. In this study, three common crops were examined, soybean, wheat and Chinese cabbage. Soybean (*Glycine max* L. Merr.) is one of the most important crop plants for seed proteins and vegetable oil [37], and wheat (*Triticum aestivum* L.) is widely planted worldwide with the caryopsis one of the staple foods for humans. Chinese cabbage (*Brassica rapa* L., *Chinensis Group*) is a dicotyledonous plant that is a popular leaf vegetable in China [38].

In this study, the capacities of *Bacillus cereus* YL6 strain to dissolve inorganic and organic P were examined. Generally, PSB dissolve insoluble-P by secreting organic acids or enzymes [39]. Based on the testing, YL6 not only produced oxalic, malonic and succinic acids but also secreted acidic, neutral, and alkaline phosphatases. The contents of the organic acids were the highest at 48 h, and similarly, the activities of the three types of phosphatases peaked at 48 h. This result indicated that 48 h was the optimum time for YL6 to dissolve insoluble P under laboratory conditions. The content of soluble-P increased rapidly in the first 48 h, which was most likely because the many acids produced and phosphatases secreted promoted PSB transformation of insoluble-P into soluble-P [40] [6]. Increased P availability improves the root growth of plants and the yield of crops. For example, Sharon isolated one efficient PSB that increased tomato growth [41]. Furthermore, growth of crops was promoted by YL6 secretions of IAA and GA, which is a result supported by other publications [42–44].

The pot experiments with soybean and wheat were conducted to determine the influences of YL6 on the growth of soybean and wheat. The numbers of PSB in rhizosphere soils of soybean and wheat increased after the addition of the YL6 strain. Therefore, PSB successfully penetrated into soil and then increased soil available P content for crop absorption. Thus, the plant total P, biomass and bean pods of soybean treated with the YL6 strain were obviously higher than those in the other treatments. In the wheat pot experiment, the content of plant total P and plant biomass with YL6 were also obviously higher than those of the groups without the YL6 strain.

The positive results of the above pot experiments resulted in determining the effects of YL6 under field conditions. Thus, YL6 was applied under field conditions to study the effects of the different treatments on the biomass and quality of Chinese cabbage. The number of soil PSB in the YL6 treatment was significantly higher than that in the other two groups in field conditions, which demonstrated that YL6 could survive in soil. Survival and colonization capacity of YL6 when inoculated in soil are the prerequisites for this PSB to play an important role in the environment and are the key factors to exert its phosphate-solubilizing capacities and help plants grow. With more of these bacteria in the soil, the conditions are more favorable for plant growth [45]. The YL6 strain also improved the growth of Chinese cabbage in field conditions.

PSB primarily rely on the ability to transform insoluble P in the soil into available P [46–48]. YL6 inoculation increased the available P in soil. Chinese cabbage could directly absorb and utilize this soil available P to promote plant growth and total P accumulation. Therefore, YL6 inoculation markedly improved growth parameters such as root and shoot dry biomass, yield and total P uptake in Chinese cabbage, compared with those of the control [46,49,50]. This result is consistent with the conclusion of Sundara, Akbari and Swarnalakshmi [49,51,52]. YL6 also increased nutritional quality indexes of Chinese cabbage, such as soluble sugar, soluble protein, and particularly vitamin C. Vitamin C is a highly active substance, which can improve human immunity and prevent cancer, heart disease and stroke but can also be used as an important contributor in fighting against aging and adversity [53]. Soluble sugar is also the material basis of polysaccharides, proteins, fats and other macromolecular compounds in plants. Our results are consistent with those of Hui [54]. Nitrate nitrogen is also an important indicator to measure vegetable quality. Because nitrite is carcinogenic and causes severe damage to the human body, improving vegetable quality by nitrate nitrogen reduction is an important task [55]. The content of nitrate nitrogen obviously decreased with the application of YL6. These results demonstrated positive effects of YL6 that improved the quality of Chinese cabbage.

The survival and colonization of PSB in the plant rhizosphere is the basis and prerequisite to promote plant growth [56]. However, in many cases, PSB do not achieve the desired effect due to insufficient numbers in the rhizosphere or failure to colonize the rhizosphere or plant [57]. In this study, the pot and field experiments showed that YL6 universally promoted the growth of the crops. Thus, the study of the colonization of the crops by YL6 is critical. GFP-labeled YL6 was used to inoculate the rhizosphere of soybean, maize and Chinese cabbage seedlings. Observation by fluorescence microscopy revealed that in seedling roots of soybean, wheat and Chinese cabbage, the GFP-labeled YL6 colonized root hairs, epidermal cells, cortex cells, intercellular spaces and vessels. The results of this study are consistent with those of other researchers [16,58–60]. Root hairs, root surfaces and epidermal cells were primarily colonized by YL6 most likely because of chemotaxis toward root exudates [61], because various carbohydrates, amino acids, organic acids and other compounds in plant root exudates are a source of nutrients for root-associated bacteria [62]. Additionally, YL6 may overcome cortex barriers by secreting cell wall degrading enzymes (CWDEs) [63]. Then, YL6 could colonize nutrient-rich intercellular spaces of plant hosts [64] and spread throughout host plants through xylem vessel lumens [65].

## Conclusions

The above experiments showed that YL6 not only dissolved soil insoluble P by secreting organic acids and phosphatases but also successfully colonized crop root tissues and promoted crop growth by secreting IAA and GA. YL6 inoculation promoted plant growth and quality and improved soil fertility. Therefore, in conclusion, the application of YL6 is a good choice for cost cutting and pollution control and to achieve high yields and reduce chemical P fertilizer use. Further research on long-term survival of PSB under field conditions and PBS colonization mechanisms is required in the future.

## Funding

The Shaanxi Science and Technology Research and Development Program (No. 2013K01-38) supported this work.

## Author Contributions

Conceptualization: YK CC.

Data curation: YK GX.

Formal analysis: GX.

Funding acquisition: CC.

Resources: FW

Software: GX

Supervision: XT CC

Validation: CC

Visualization: YK XY

Writing – original draft: YK GX

Writing – review & editing: YK GX HL

